# Robust Hierarchical Co-clustering to Explore Toxicogenomic Biomarkers and Their Regulatory Doses of Chemical Compounds

**DOI:** 10.1101/2020.05.13.094946

**Authors:** Mohammad Nazmol Hasan, Md. Bahadur Badsha, Md. Nurul Haque Mollah

## Abstract

Toxicogenomics combines high throughput molecular technologies with statistical and machine learning approaches to discover a similar group of doses of chemical compounds (DCCs) and genes to explore toxicogenomic biomarkers and their regulatory DCCs. This is also very important in the toxicity study of environmental stressors, synthetic chemicals and drug discovery and development process. Different clustering algorithms are concerned with the discovering of interesting clusters/groups of row or column entities of a dataset. Among those hierarchical clustering (HC) and logistic probabilistic hidden variable model (LPHVM) can identify toxicogenomic biomarkers and their regulatory DCCs forming co-cluster. However, the HC method is very sensitive to outlying observations. On the other hand, though LPHVM is a robust approach, it consumes more time for calculation since it is Expectation-Maximization (EM) based iterative approach. Additionally, the LPHVM creates artificiality problem taking absolute value of the data matrix. Therefore, to overcome these problems in this paper, we proposed a robust hierarchical co-clustering (RHCOC) algorithm to co-cluster genes and DCCs simultaneously with a view to explore toxicogenomic biomarkers and their regulatory DCCs. The performance of the proposed RHCOC algorithm over the conventional HC for clustering genes and DCCs of toxicogenomic data has been investigated based on the simulation study. The results of the simulation study have shown that the RHCOC approaches produce far lower clustering error rate (ER) than the conventional HC approaches in presence of outlying observations in the dataset. Otherwise they perform equally in absence of outlier in the dataset. To explore biomarker co-clusters consisting of toxicogenomic biomarker genes and their regulatory DCCs we used control chart for individual measurement (CCIM). We have also investigated the performance of the proposed approach in the case of the pathway level real life fold change gene expression (FCGE) toxicogenomic data analysis. The biomarker co-clusters consisting of toxicogenomic biomarker genes and their regulatory DCCs and biomarker genes explored by the proposed approaches have been validated by the literature and functional annotation. Our method is implemented in R package “rhcoclust” available on github (https://github.com/mdbahadur/rhcoclust).

## 1 INTRODUCTION

Toxicogenomics examines how the entire genetics structure is involved in an organism’s responses to chemical agents, including drugs and environment stressors. It received increasing attention in genetics with the rapid advancement of high throughput molecular profiling technologies such as transcriptomics, proteomics and metabolomics (Hamadeh et al., 2002; Waters and Fostel, 2004; Ancizar-Aristizábal et al., 2014). Statistical algorithms or machine learning techniques together with these high throughput technologies can identify toxicogenomic biomarkers as well as asses chemicals’/drugs’ safety (Chung et al., 2015; Hasan et al., 2018). The toxicogenomic biomarkers are those genes or proteins or metabolites which become up/down-regulated exposure to chemicals/drugs. On the other hand, from the concept of central dogma ‘the expression of a gene regulates the expression of its products (proteins and metabolites)’ (Zhu et al., 2005). Therefore, chemicals/drugs toxicity can be observed from the gene expression analysis of target organs before the phenotypic change has been appeared (Fielden et al., 2007; Uhara et al., 2008; Igarasi et al., 2015).

In toxicogenomic studies and in drug discovery and development process it is very important to know doses of chemical compounds (DCCs)/drugs similarity or groups according to the mechanism of action as well as groups of genes which have a similar expression patterns in response to these DCCs groups (Madeira and Oliveira, 2004; Afshari et al., 2011; Hasan et al., 2018). The clustering algorithms are concerned with the discovery of interesting groups of row or column entities of the data matrix with large heterogeneity among them of a dataset based on the values of these entities. Different clustering algorithms are developed for a variety of application domain which has been proven to be very effective (Ward Jr, 1963; MacQueen, 1967; Bezdek, 1981; Kohonen, 1989). Some of these algorithms have been and are presently being implementing to retrieve the latent patterns in gene expression data (Eisen et al., 1998; Tavazoie et al., 1999; Tamayo et al., 1999; Chung et al., 2015; Hasan et al., 2018; Hasan et al., 2019). Although, some success observed with the existing clustering algorithms in gene expression data analysis, still not a single clustering algorithm is developed, which can be the most suitable for gene expression data (Eisen et al., 1998; Tavazoie et al., 1999; Tamayo et al., 1999; Ben-Dor et al., 1999; Quackenbush, 2001; Herrero, 2001). Furthermore, most of the clustering algorithms likewise K-means, model based, self-organizing map (SOM), fuzzy, probabilistic hidden variable model (PHVM) cluster the objects based on the predetermined number of clusters in the dataset. Nonetheless, the cluster selection methods produce different results for the same dataset. The hierarchical clustering (HC) algorithm visualizes the clusters as a tree-like dendrogram and correlations between genes or DCCs. The researcher can choose the number of clusters for the dataset observing the dendrogram in a convenient way. But, the HC algorithm offers different combinations of distance and clustering methods for grouping objects. Nevertheless, all of them are not equally suitable for clustering genes and DCCs of toxicogenomic data. Recently, a method has developed that the combinations of distance and clustering methods euclidean: ward, manhattan: ward and minkowski: ward are suitable for clustering as well as co-clustering (clustering row (gene) and column (DCCs) entities of a data matrix simultaneously) genes and DCCs (Hasan et al., 2019b). However, the algorithm has two important shortfalls. Firstly, it does not identify the biomarker co-clusters with statistical justification. Secondly, the search of groups/clusters or co-clusters based on these combinations is completely spoiled if there are outlying observations that exist in the dataset. On the other hand, microarray based toxicogenomic data often contains outlying observations. These outlying observation occurs due to various types of noises are normally arises at different experimental stages during data collection. For example, when producing the DNA array, preparing biological samples, and extracting of the results, etc. (Berrar et al., 2003; Gottardo et al., 2006; Upton et al., 2009). Though, logistic probabilistic hidden variable model (LPHVM) (Hasan et al., 2018) can cluster as well as co-cluster the toxicogenomic data based on the predetermined number of clusters in absence and presence of outliers in the dataset, but the problem is it consumes more time for computation. Because, LPHVM is an Expectation-Maximization (EM) algorithm (Dempster et al., 1977) based iterative approach. Furthermore, the LPHVM creates artificiality considering absolute value of the dataset. This artificiality grouped the up-regulated and down-regulated gene sets into a single cluster which response to a set of DCCs of a similar mechanism of action. Though, practically these sets of genes are the member of two different clusters. Therefore, to overcome these problems, we proposed robust hierarchical co-clustering (RHCOC) algorithm and control chart for individual measurement (CCIM). The RHCOC is the robust approach for clustering as well as co-clustering genes and DCCs in the absence and presence of outlying observations in the dataset. The CCIM method identify the biomarker co-clusters (co-clusters consisting of toxicogenomic biomarker genes and their regulatory DCCs) to explore toxicogenomic biomarkers and their regulatory DCCs based on the RHCOC algorithm results.

## 2 METHODS AND MATERIALS

### 2.1 Description of Toxicogenomic Data

In toxicogenomic studies the experiments are set out to measure the gene expression of treatment and control group of animal samples. The treatment group of animal samples are treated with doses of chemical compounds (DCCs) at multiple time points and control group of animals are kept free of treatments. **Figure 1** represented a usual toxicogenomic experiment and gene expression data from the treatment and control group of animal samples are collected. The fold change gene expression (FCGE) data which computed from this experiment using the equations (1) and (2) are used in most of the toxicogenomics studies. Because FCGE dataset directly reflects the treatment effects (Nyström-Persson et al., 2013; Chung et al., 2015; Nyström-Persson et al., 2017; Hasan et al., 2018, 2019). The fold change gene expression *U_pqrt_* for the *p^th^* (*p* = 1, 2,⋯*P*) chemical compound, *q^th^* (*q* = 1,2,3) dose level, *t^th^* (*t* = 1,2,⋯,*T*) time point and *r^th^*(*r* = ι, 2, 3) animal sample and *U_pqr_* for single time point can be computed from the gene expression data of this experiment using the equations:

**Figure 1.**
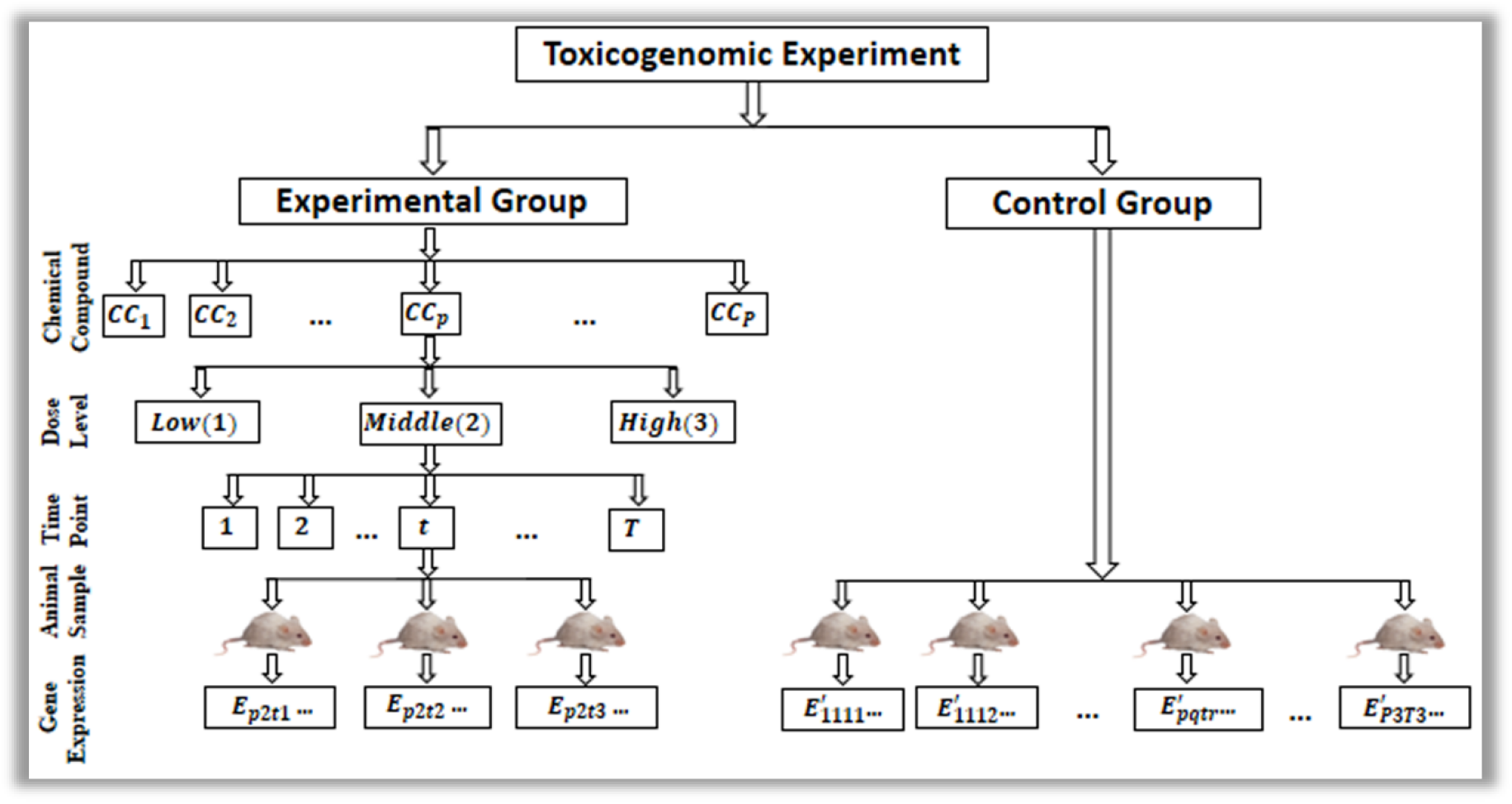
A usual toxicogenomic experiment.

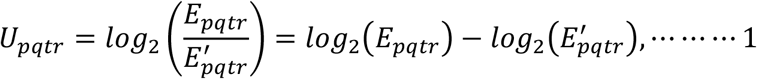

and

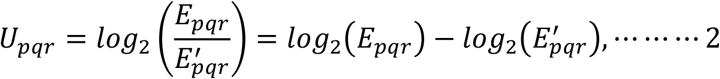

respectively. The average FCGE value over the animal samples of a gene are *Ū_pqt_*. and *Ū_pq_* respectively for multiple and single time point. From these values the effect of DCCs over the genes can be measured. Average FCGE value will be positive for up-regulated genes and negative for the down-regulated genes. In the literature it is proved that the combinations euclidean: ward, manhattan: ward and minkowski: ward of distance and HC methods are suitable for clustering or co-clustering genes and DCCs of toxicogenomic data. However, these conventional combinations cannot cluster/co-cluster genes and DCCs properly in the presence of outlying observations in the dataset. Thus, in the following sections we discuss about the robust procedure of conventional HC approaches.

### 2.2 Logistic Transformation of Fold Change Toxicogenomics Data

Transformation based outlier reducing approaches are more popular and comparatively convenient for the users to obtain robust results from the methods (Atkinson, 1982; Carroll, 1982; Box and Cox, 1964, Hasan et al., 2019a). Among all the transformation based approaches logistic transformation based approach is more efficient to obtain robust results (Hasan et al., 2018). Before application of logistic transformation in the dataset we have taken average value *Ū_pqt_* of the fold change gene expression (*U_pqtr_*) over the animal samples. Then, applying logistic transformation on the average FCGE value (*Ū_pqt_*), we get transformed *n* × *m* (gene-DCCs) fold change gene expression data matrix consisting of G = (*G*_1_, *G*_2_,…, *G_n_*) genes and C = (*C*_1_, *C*_2_, …, *C_m_*) DCCs. We termed each of the input of the transformed data matrix as *Tr*(*G_i_, C_j_*) for the convenience of further use which represents the transformed FCGE value for the *i^th^* gene under the *j^th^* treatment conditions. Where *Tr*(*G_i_, C_j_*)can be expressed as:

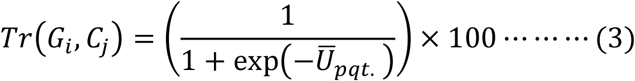

### 2.3 Robust Hierarchical Co-clustering (RHCOC) Algorithm

In our proposed RHCOC algorithm we shall apply the mentioned conventional combinations of distance and HC approaches (euclidean: ward.D, euclidean: ward.D2, manhattan: ward.D, manhattan: ward.D2, minkowski: ward.D or minkowski: ward.D2) on the logistically transformed FCGE data for robust clustering and co-clustering genes and DCCs. To co-cluster genes and DCCs the number of clusters in the genes (row entities) and DCCs (column entities) are estimated observing the dendrogram (hierarchical clustering) of the data matrix. The reasons behind using dendrogram for this purpose described in section 1 (introduction). The average FCGE value are computed for every intersection areas of the gene and DCCs clusters generated by the HC method. The gene and DCCs clusters together with make co-cluster. The biomarker co-clusters (co-clusters consisting of toxicogenomic biomarkers and their regulatory DCCs) can be discovered using the control chart for individual measurement (CCIM). Then from these biomarker co-clusters the toxicogenomic biomarkers and their regulatory DCCs are explored. These findings are very important for the biologists, drug developers and the researchers in the field of toxicogenomics. The detail description of the proposed RHCOC algorithm for the exploration of toxicogenomic biomarkers and their regulatory DCCs are given bellow.

**Step 1:** The outlier effects in the dataset are reduced using the logistic function 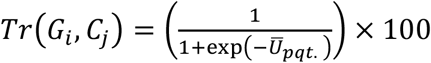 where *Ū_pqt._*. is the observation is and *Tr*(*G_i_, C_j_*) is the transformed value of that observation for *i^th^* row and *j^th^* column of the data matrix which is expressed in %. For this study we consider *Tr*(*G_i_, C_j_*) is the transformed value of the fold change gene expression (FCGE) value of the *i^th^* gene under the *j^th^* treatment conditions (*p^th^* chemical compound, *q^th^* dose level, *t^th^* time point).
**Step 2:** The number of cluster in the genes (row entity) and in the doses of chemical compounds (DCCs)/treatment conditions (column entity) of the data matrix are estimated observing the dendrogram. The fixation of the number of clusters for the row and column entities depends on the researchers’ interest.
**Step 3:** Rearrange the transformed data matrix according to the row (gene) and column (DCCs) clusters generated by the hierarchical clustering.
**Step 4:** Each of the row (gene) cluster has an intersection area with each of the column (DCCs) cluster in the rearranged data matrix generated in **step 3**. We take the average of the FCGE value of the intersection areas for every pair of the row and column clusters. For example, if there are four row clusters and three column clusters, then there will be 12 pair (combinations) of row and column clusters. Each pair/combination of the row and column cluster makes co-cluster themselves. We term the average FCGE value in the intersection area as the co-cluster mean.
**Step 5:** Rank the average FCGE values of the co-clusters (computed in **step 4**) in descending order and their associated pair/combination of row and column clusters rearranged accordingly.
**Step 6:** We assign cluster number for row (genes) and column (DCCs) clusters according to the ranked average FCGE values which we get from **step 5**. The co-cluster having largest average FCGE value is considered as the co-cluster 1. The row cluster and the column cluster that make the co-cluster 1 are also termed as row/gene cluster 1 and column/DCCs cluster 1. Similarly, the co-cluster having the second largest average FCGE value is termed as co-cluster 2 and row/gene and column/DCCs cluster associated with co-cluster 2 are termed as row/gene cluster 2 and column/DCCs cluster 2 and so on.
**Step 7:** Finally we explore biomarker co-clusters using control chart for individual measurement (CCIM). The description of the CCIM are given in section **S3** of the supplementary material and in section **2.4** of the main article. The co-clusters which average FCGE value plots beyond the UCL and LCL is termed as biomarker co-clusters. The genes and DCCs that consist in the biomarker co-clusters are termed as the toxicogenomic biomarker genes and their regulatory DCCs and we extract them from the biomarker co-clusters.

### 2.4 Extraction of Toxicogenomic Biomarkers and Their Regulatory DCCs

As mentioned earlier that the toxicogenomic biomarkers are those genes or proteins or metabolites which become up/down-regulated exposure to environmental chemical or synthetic chemical compounds/drugs. Therefore, according to RHCOC algorithm described in the previous section the cluster of up-regulated biomarker genes makes co-cluster with their regulatory DCCs cluster with larger average FCGE values in their intersection area. Similarly, the down-regulated biomarker gene cluster makes co-cluster with their regulatory DCCs cluster with the smaller average FCGE values in their intersection area. We term these co-clusters as the biomarker co-clusters. These co-clusters have assignable variations from the other co-clusters of equally expressed genes and non-regulatory DCCs. To separate these biomarker co-clusters from the non-biomarker co-clusters we use CCIM. The CCIM is a special type of control chart is preferable for individual measurement or when the sample size is one for observing the quality characteristics of a product in a production process. This control chart is efficient to differentiate assailable causes of variation (variations due to improperly control machine, operators’ errors, and defective raw materials) from the chance/natural causes of variation of a production process (Montgomery, 2016, Hasan et al., 2019a). The CCIM contains a central line (CL) which represents the average value of the quality characteristics corresponding to in-control state, an upper control limit (UCL) and a lower control limit (LCL). If any observation plots outside the control limits (UCL or LCL) it indicates that the process running with assignable causes of variation action is needed to control it. On the other hand, any observation plotting within the control limits (UCL and LCL) it indicates is that the process running with natural causes of variation and action is not needed. Wheeler, 1995 has said that the control charts work irrespective of the distribution, so CCIM can be used for individual measurement even the observations do not follow normal distribution. In fact, there is an analogy between the process monitoring consisting of individual measurement and assessing the DCCs/drugs toxicity based on toxicogenomic data (Hasan et al., 2019a). Therefore, in this study, we have used CCIM to identify the biomarker co-clusters to explore toxicogenomic biomarkers and their regulatory DCCs. The description of CCIM are given in the supplementary section S3.

### 2.5 Robustness of the Proposed RHCOC Algorithm

There are three ways of getting robust outputs in presence of outlying observations in the dataset: i) applying the robust methods ii) deleting the outlying observations and iii) applying conventional methods on the transformed dataset. The first approach is more complicated than using the classical methods for the users. On the other side, deletion of outlying observation loses the information of dataset. Thus, the last approach transformation based robust methods is more popular and convenient for the users to obtain robust results from the methods in presence of outlying observations in the dataset (Atkinson, 1982; Carroll, 1982; Box and Cox, 1964, Hasan et al., 2019a). Therefore, in this study we consider logistic transformation for reducing outlier effects from the dataset. Because, if there are extreme values in the dataset the logistic transformation bring them within the range of 0–1. The other transformation methods like Box-Cox family of power transformation returns unbounded value for the extreme one (Hasan et al., 2018). Therefore, logistic transformation based proposed RHCOC algorithm produce more robust results compare to the other methods.

### 2.6 Clustering Performance Measurement

We investigate the performance of the RHCOC approach with a conventional HC approach using simulated datasets in the absence and presence of outliers in the dataset based on the clustering error rate (ER). The genes and DCCs which are considered in one cluster in the simulated dataset are incorrectly assigned in another cluster by the RHC or HC is considered as the miss clustered observations. We define ER as the percentage of miss clustered observations which is calculated as:

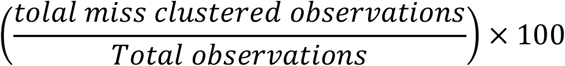

### 2.7 Simulated Dataset

To investigate the performance of the conventional distance and HC method combinations (euclidean: ward.D, euclidean: ward.D2, manhattan: ward.D, manhattan: ward.D2, minkowski: ward.D and minkowski: ward.D2) with our proposed RHCOC approaches for clustering genes and DCCs in absence and presence of outlying observations we use simulated dataset. In the simulated dataset we construct a pathway level FCGE data of size (*n* = 50 × *m* = 36). We consider the simulated dataset at 24 hour time point since toxicity effect visualize more clearly at this time point (Nyström-Persson et al., 2013). The characteristic of toxicogenomic gene expression data is that some genes are up-regulated and some are down-regulated due to the exposure of same set of DCCs. We generated FCGE data (*U_pqr_*) according to the characteristics of toxicogenomic data using the following model:

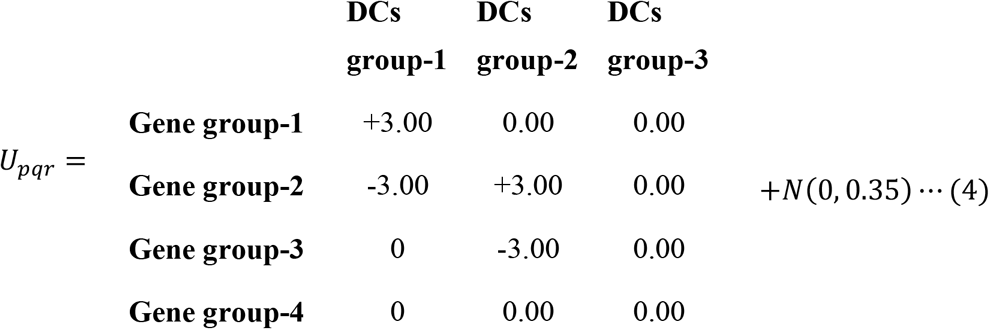

There are four gene groups and three DCCs groups in the simulated dataset. The gene group 1, 2, 3 and 4 consists of genes *G*1 – *G*10, *G*11 – *G*20, *G*21 – *G*30 and *G*31 – *G*50 respectively and DCCs group 1, 2 and 3 consists the DCCs C1_High-C5_High-C1_Middle-C5_Middle, C6_High-C10_High-C6_Middle-C10_Middle and C1_Low-C12_Low-C11_Middle-C12_Middle-C11_High-C12_High respectively. Where, G stands for gene and C stands for doses of chemical compound are arranged in the row and column of the simulated data matrix respectively. The error term *N*(0, 0.35) from normal distribution with mean 0 and variance 0.35 is added to each element of the simulated dataset. In the simulated dataset the gene group-1 is up-regulated by the DCCs group-1, gene goup-2 is up and down-regulated by the DCCs group-2 and 1 respectively. The gene group-3 is down-regulated by the DCs group-2. The gene group-4 is not regulated by any of the DCCs groups and DCCs group-3 does not influence any of the genes in the dataset.

### 2.8 Simulated Data Contamination by Outliers

An outlier is an observation in the dataset that arises for some unexpected circumstances which deviate from the actual value of the observation. It is a common problem for gene expression/fold change gene expression data analysis. To compare the performance of the RHCOC approach with conventional HC approaches in the absence or presence of outlying observations in the dataset, we contaminate the simulated FCGE dataset inputting outliers in the dataset. The outlying observations in the simulated dataset are defined as the five times larger FCGE values than the maximum value in the original dataset. These outlying observations may arise in the dataset casewise following the Tukey-Huber contamination model (THCM) (Agostinelli et al., 2015; Hasan et al., 2018) or independent cellwise following the independent contamination model (ICM) (Alqallaf et al., 2009; Hasan et al., 2018). In our gene-DCCs FCGE toxicogenomic data matrix each gene in the row is a case and each input in the data matrix is a cell. Therefore, for investigating the robustness of the proposed RHCOC approaches over the conventional HC approaches we have contaminated the simulated dataset by outliers casewise and independent cellwise. The descriptions of casewise or THCM and independent cellwise or ICM data contamination methods are given in the S2 (S2.1 and S2.2).

### 2.9 Real Datasets

To show the application and performance of the proposed RHCOC algorithm in the real aspect we use pathway level gene expression data of the Japanese Toxicogenomics Project (TGP) (Uehara et al., 2010). Here we consider pathway level FCGE data from *Rattus Norvegicus, in vivo*, liver, multiple dose experiment at 4 time points (3 hours, 6 hours, 9 hours, 24 hours). We have downloaded FCGE data of the glutathione metabolism pathway (GMP) and PPAR signaling pathway (PPAR-SP) from “Toxygates” (https://toxygates.nibiohn.go.jp/toxygates/#columns) (Nyström-Persson et al., 2013) which is an online toxicogenomic data analysis as well as a database platform. In case of the GMP dataset, we consider all dose levels (Low, Middle and High) of three glutathione depleting compounds (acetaminophen, methapyrilene and nitrofurazone) and seven non-glutathione depleting compounds (erythromycin, hexachlorobenzene, isoniazid, gentamicin, glibenclamide, penicillamine and perhexilline) (Nyström-Persson et al., 2013). On the other side, for PPAR-SP dataset we consider PPARs related gene regulatory compounds (WY-14643, clofibrate, gemfibrozil, benzbromarone and aspirin) (Kiyosawa et al., 2006) and some other randomly selected compounds (cisplatin, diltiazem, methapyrilene, phenobarbital and triazolam) along with their dose levels low, middle and high at multiple time points.

## 3 RESULTS

### 3.1 Simulation Study

It is mentioned earlier that the combinations euclidean: ward, manhattan: ward and minkowski: ward of distance and HC methods are suitable for clustering/co-clustering toxicogenomic data (Hasan et al., 2019b). In this paper, we have also claimed that these combinations of distance and clustering methods are very sensitive to outlying observations in the dataset. In our proposed RHCOC algorithm we have applied the combinations of distance and clustering method (euclidean: ward.D, euclidean: ward.D2, manhattan: ward.D, manhattan: ward.D2, minkowski: ward.D and minkowski: ward.D2) on the logistically transformed gene-DCC FCGE data matrix. Here we have used the methods ‘word.D’ and ‘word.D2’ instead of the ‘word’ since there are two approaches (word.D and word.D2) of word method. In the **Figure 2**, we have presented the ER for clustering genes and DCCs in the absence and presence of outlying observations in the simulated dataset. We have also depicted the average clustering ER (over the rate of outliers; THCM: 0%-40%, ICM: 0.00-0.598) against the combinations of distance and HC methods for the classical and proposed approaches **Figure S3** (supplementary material). Both the classical and the proposed approaches produce 0% ER (perform equally) in the absence of outlying observations in the simulated dataset. Nevertheless, in the case of outliers in the dataset it is observed that the proposed RHCOC approach outperforms the classical approach (**Figure 2 and Figure S3 (supplementary material))**. Observing all the plots given in **Figure 2** and in the supplementary material we can conclude that the distance and HC method combination manhattan:ward.D produces lower ER, as well as more stable results, compare to the others proposed approaches (euclidean:ward.D, euclidean:ward.D2, manhattan:ward.D2, minkowski:ward.D and minkowski:ward.D2) for clustering genes or DCCs is considered as the best approach for toxicogenomic data. Co-clustering (clustering genes and DCCs simultaneously) genes and DCCs is one of the important objectives of toxicogenomic studies as well as drug development processes (Afshari et al., 2011; Hasan et al., 2018; Hasan et al., 2019b). Therefore, we have proposed the RHCOC algorithm for co-clustering genes and DCCs to explore toxicogenomic biomarkers and their regulatory DCCs. According to this algorithm, we assign the gene cluster as well as DCCs cluster as 1 which produces the largest co-cluster mean. Similarly, we assign the gene cluster as well as DCCs cluster as 2 which produce the second largest co-cluster mean and so on. Dendrogram to detect the number of clusters in the gene (row entity) and in the DCC (column entity) and the co-cluster graph are given in the supplementary figures **Figure S1** and **Figure S4** respectively. The RHCOC algorithm arranges gene and DCCs clusters according to the ranked co-clusters mean (for detail see section **2.3**). The ranked co-cluster mean and their associated gene and DCC clusters for the simulated data are given in **Table 1**. Furthermore, for visualizing the performance of the proposed approaches over the classical approaches the original (simulated) co-cluster structure, gene and DCCs randomly allocated structure and the classical and proposed approach reconstructed structure in absence and presence of outlying observations (10%) are given in **Figure 3**. From these information it is observed that the proposed method produces lower ER and reconstructs the original simulated data structures more efficiently than the classical approaches. We have identified the biomarker co-clusters applying the control chart for individual measurement (CCIM) (described in section 2.4 and **S3** (supplementary material)) on the RHCOC algorithm generated co-clusters mean and their respective gene and DCC clusters. Where we termed the biomarker co-cluster as the co-cluster consisting of biomarker genes and their regulatory DCCs. The biomarker co-clusters together with the respective gene and DCC clusters for the simulated data are given in the **Figure S4** (supplementary material).

**Figure 2.**
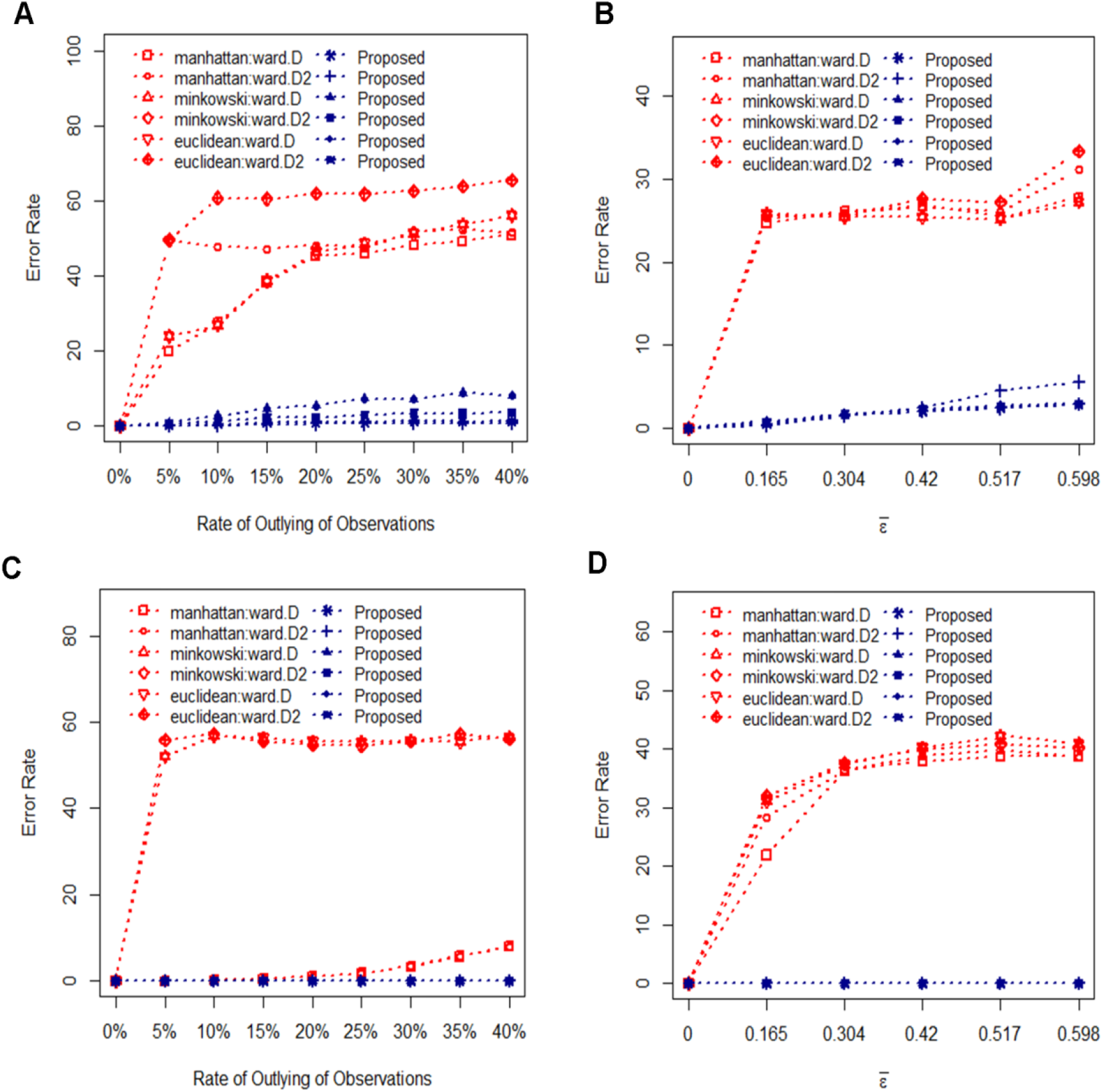
The percentage of error rate is plotted against the rate of outlying observations when the dataset is simulated 100 times. (**A**) Gene clustering ER in case of THCM data contamination method. (**B**) Gene clustering ER in case of ICM data contamination method. (**C**) DCCs clustering ER in case of THCM data contamination method. (**D**) DCCs clustering ER in case of ICM data contamination method.

**Figure 3.**
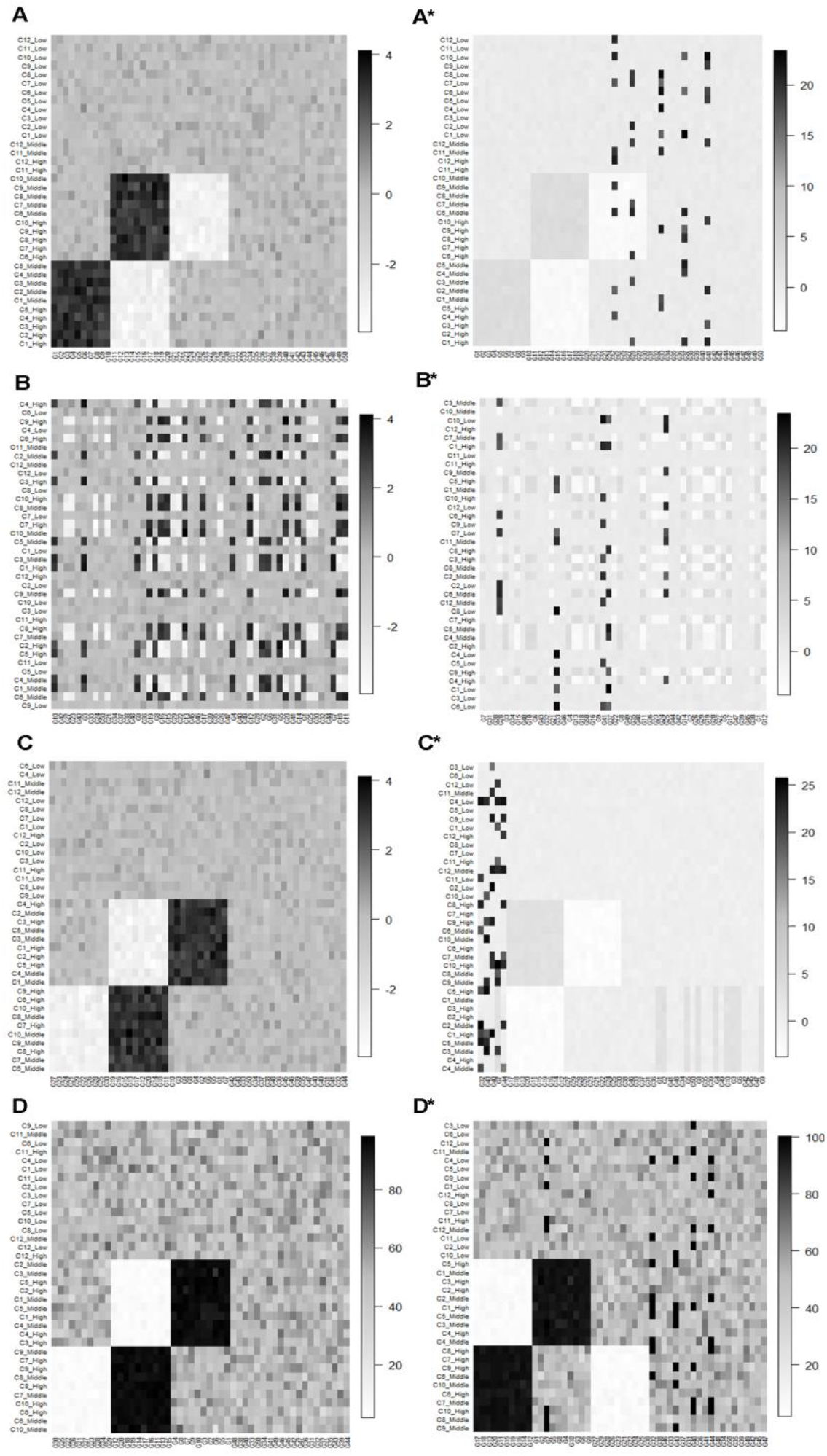
A view of the classical and proposed RHCOC algorithm reconstructed data structures in the absence and presence (10%) of outliers. **(A)** The simulated data structure in the absence, and **(A*)** in the presence of outliers. (B) The randomly allocated data structure in the absence, and (B*) in the presence of outliers. **(C)** Classical method reconstructed data structure in the absence, and **(C*)** in the presence of outliers. **(D)** RHCOC algorithm reconstructed data structure in the absence, and **(D*)** in the presence of outliers.

**Table 1.** The average ranked co-cluster means along with the gene and DCC cluster numbers for the simulated and real datasets generated by the RHCOC algorithm. In the first column (**Cluster #**) the first and second number represents the gene and DCC cluster number respectively.

| Simulated Data |  | GMP Data |  | PPAR-SP Data |  |
| --- | --- | --- | --- | --- | --- |
| Cluster # | Co-Cluster Mean | Cluster # | Co-Cluster Mean | Cluster # | Co-Cluster Mean |
| 1,1 | 94.912 | 1,1 | 70.434 | 1,1 | 79.117 |
| 2,2 | 94.910 | 1,2 | 51.229 | 1,2 | 62.867 |
| 1,3 | 50.726 | 2,1 | 50.387 | 2,1 | 53.373 |
| 3,1 | 50.208 | 3,1 | 50.084 | 2,2 | 49.663 |
| 4,3 | 49.917 | 2,2 | 49.819 | 2,3 | 47.837 |
| 2,3 | 49.911 | 2,3 | 47.828 | 1,3 | 47.601 |
| 3,3 | 49.849 | 3,2 | 47.299 | 3,2 | 46.600 |
| 4,2 | 49.680 | 1,3 | 46.344 | 3,3 | 46.470 |
| 2,1 | 49.570 | 3,3 | 43.229 | 3,1 | 45.823 |
| 3,2 | 49.501 |  |  |  |  |
| 1,2 | 4.833 |  |  |  |  |
| 4,1 | 4.787 |  |  |  |  |

### 3.2 Real Data Analysis

To investigate the performance of the proposed RHCOC algorithm in the case of real life toxicogenomic data analysis we consider glutathione metabolism pathway (GMP) and PPAR signaling pathway (PPAR-SP) datasets (described in section 2.5) for multiple time points (3 hours, 6 hours, 9 hours, 24 hours) and Low, Middle and High dose levels. In case of the GMP dataset we consider three glutathione depleting compounds (acetaminophen, methapyrilene and nitrofurazone) and seven non-glutathione depleting compounds (erythromycin, hexachlorobenzene, isoniazid, gentamicin, glibenclamide, penicillamine and perhexilline) (Nyström-Persson et al., 2013) and for PPAR-SP dataset we consider PPARs related gene regulatory compounds (WY-14643, clofibrate, gemfibrozil, benzbromarone and aspirin) (Kiyosawa et al., 2006) and some other randomly selected compounds (cisplatin, diltiazem, methapyrilene, phenobarbital and triazolam). The gene and DCCs clustering (dendrogram) considering all dose levels and multiple time points for GMP and PPAR-SP datasets are given in **Figure S2** (supplementary material). Observing these dendrograms we decide there are three clusters in the genes and DCCs for both of the GMP and PPAR-SP datasets. Then ranked means of the gene and DCC co-clusters along with their respective cluster numbers for the genes and DCCs are given in **Table 1**. According to the RHCOC algorithm, the clusters of genes and DCCs are given in **Table 2** (GMP dataset) in **Table S2** (PPAR-SP datasets) **in supplementary material**. Gene and DCCs clusters for the PPAR-SP and simulated datasets are given in the supplementary material. The co-cluster view based on the RHCOC algorithm for the real datasets (GMP and PPAR-SP) are depicted in **Figure 4**. The same view of the simulated dataset is given in the supplementary material (**Figure S4**). The biomarker co-clusters along with the respective gene and DCCs clusters are explored applying the CCIM method on the RHCOC algorithm generated results. The graphical view of the CCIM method for the simulated, GMP and PPAR-SP datasets are given in **Figure S5** (supplementary material). We extract the toxicogenomic biomarker genes and their regulatory DCCs from the biomarker co-clusters. The biomarker genes for the GMP and PPAR-SP datasets are functionally annotated via DAVID (Huang da et al., 2009) and found highly significant in the respective pathways which are given in **Table 3** (GMP) and **Table S3** (supplementary materials) (PPAR-SP).

**Figure 4.**
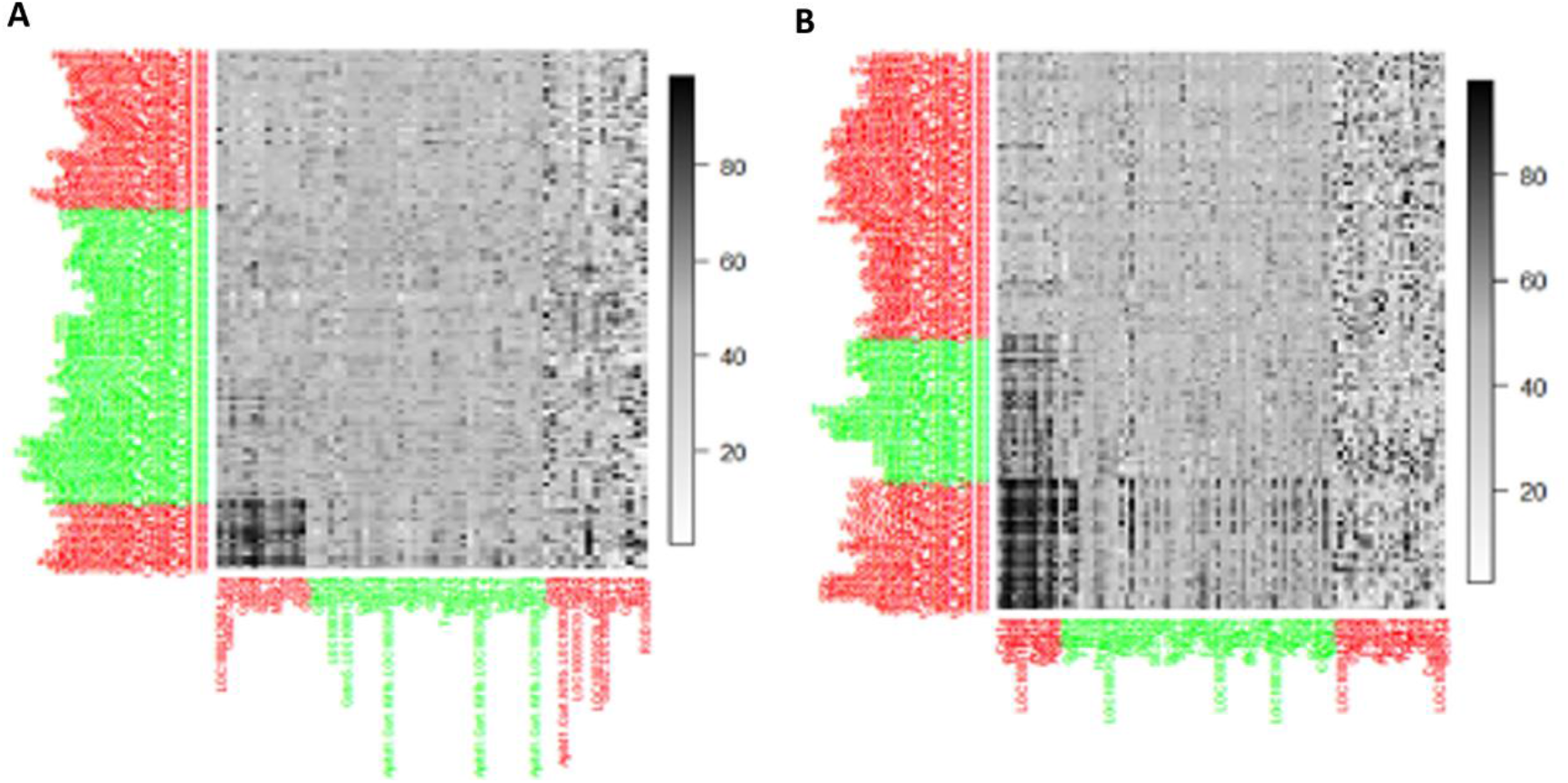
The proposed RHCOC algorithm recovered co-clustered data structure. **(A)** GMP data. **(B)** PPAR-SP data.

**Table 2.**
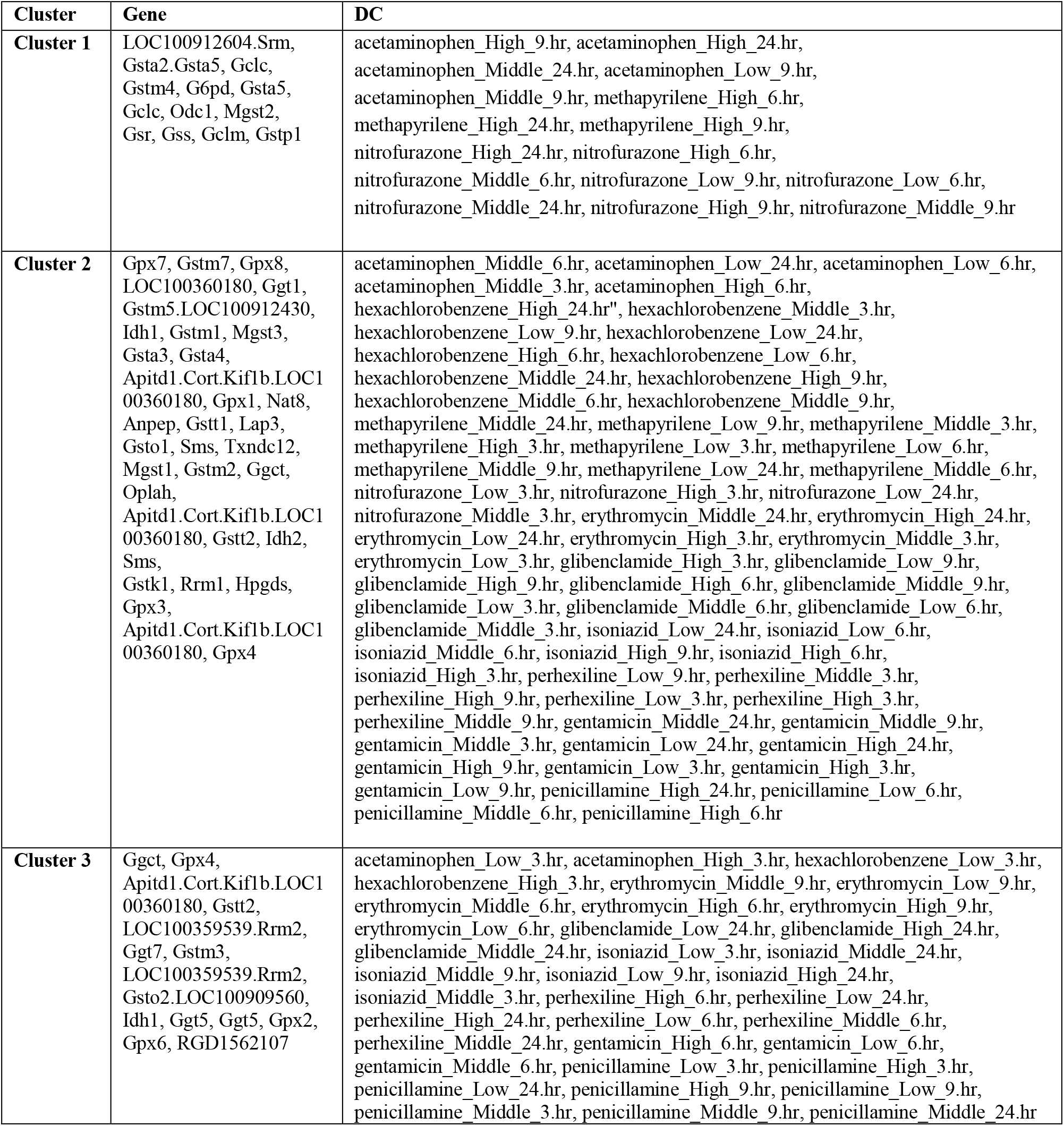
The gene and DCCs clusters of the GMP dataset retrieve by the proposed RHCOC algorithm.

**Table 3.** Functional annotation of the KEGG pathway on the biomarker genes (genes in cluster 1) explored by the RHCOC algorithm for glutathione metabolism pathway datasets.

| Term (KEGG pathway) | Count | % | p-value | Genes |
| --- | --- | --- | --- | --- |
| rno00480:Glutathione metabolism | 11 | 91.66 | 2.43E-22 | Mgst2, G6pd, Gclm, Odc1, Gsr, Gsta5, Gclc, Gclc, Gsta2/Gsta5, Gstp1, Gss, Gstm4 |
| rno00980:Metabolism of xenobiotics by cytochrome | 5 | 41.66 | 1.23E-6 | Mgst2, Gsta5, Gsta2/Gsta5, Gstp1, Gstm4 |
| rno00982:Drug metabolism - cytochrome | 5 | 41.66 | 1.30E-6 | Mgst2, Gsta5, Gsta2/Gsta5, Gstp1, Gstm4 |
| rno05204:Chemical carcinogenesis | 5 | 41.66 | 3.54E-6 | Mgst2, Gsta5, Gsta2/Gsta5, Gstp1, Gstm4 |
| rno01100:Metabolic pathways | 5 | 41.66 | 0.068 | G6pd, Gclm, Odc1, Gclc, Gclc, Gss |

## 4 DISCUSSION AND CONCLUSIONS

Many authors’ consensus is that identification of subsets of DCCs which have similar mechanism of action over the respective subsets of genes as well as identification of the biomarker genes and their regulatory DCCs are the main objectives of toxicogenomic studies as well as drug development process (Madeira and Oliveira, 2004; Afshari et al., 2011; Nyström-Persson et al., 2017; Hasan et al., 2018; Hasan et al., 2019b). The HC algorithm is more popular and widely used method which can be used for clustering genes and DCCs. Nonetheless, this algorithm does not make co-clusters between genes and DCCs. PHVM and LPHVM co-cluster the objects based on the predetermined number of clusters in the dataset (Zhu et al., 2005; Joung et al., 2006; Bicego et al., 2010; Chung et al., 2015; Hasan et al., 2018). Nonetheless, these methods consumes more time for computation since they are Expectation-Maximization (EM) algorithm (Dempster et al., 1977) based iterative approach. Determination of number clusters in the dataset is also ambiguous since different cluster selection methods produce different results for the same dataset. Furthermore, the LPHVM creates artificiality considering absolute value of the dataset. In this aspect Hasan et al., 2019b proposed a co-clustering algorithm and showed that the algorithm co-cluster the genes and DCCs efficiently to explore toxicogenomic biomarkers and their regulatory DCCs. The author also proved that the combinations of distance HC methods (euclidean: ward, manhattan: ward and minkowski: ward are suitable combinations for clustering genes and DCCs of toxicogenomic data. However, all of these approaches are very sensitive to outlying observations. On the other side, microarray based toxicogenomic data often contains outlying observations. Because, various types of noise are normally arises at different experimental stages during data collection. For example, when producing the DNA array, preparing biological samples, and extracting of the results, etc. (Berrar et al., 2003; Gottardo et al., 2006; Upton et al., 2009). To overcome these problems in this paper, we have proposed robust hierarchical co-clustering (RHCOC) algorithm to cluster as well as co-cluster genes and DCCs. In the case of RHCOC, we have applied the distance and HC method combinations (euclidean: ward.D, euclidean: ward.D2, manhattan: ward.D, manhattan: ward.D2, minkowski: ward.D and minkowski: ward.D2) on the logistically transformed fold change gene expression (FCGE) data. Since logistic transformation brings the outlying observations within the reasonable space without changing the patterns of genes and DCCs in the dataset. Hence RHC produces robust results.

The performance of the proposed RHCOC over the conventional HC for clustering genes and DCCs of toxicogenomic data has been investigated based on the simulation study. The results of the simulation study have shown that the RHCOC approaches produce far lower clustering error rate (ER) than the conventional HC approaches in presence of outlying observations otherwise they perform equally in absence of outlier in the dataset. Therefore, RHCOC approaches outperform over the conventional HC approaches. We also have investigated the performance of the proposed algorithm in the case of the pathway level real life FCGE toxicogenomic data analysis. The biomarker co-clusters produced by the RHCOC algorithm have been validated with the literature and functionally annotated. The toxicogenomic biomarker genes explored by the biomarker co-clusters are highly statistically significant in the respective pathways and DCCs having the toxicity evidence (,Kiyosawa *et al*., 2006; Huang da et al., 2009; Nyström-Persson et al., 2013).

## Supporting information

https://bsmrau.edu.bd/nazmol/other-activities/

## 5 DATA AVAILABILITY

The real dataset is available at https://toxygates.nibiohn.go.jp/toxygates/#columns

## 6 AUTHOR CONTRIBUTION

The idea or concept of the RHCOC algorithm and R programming code developed by Mohammad Nazmol Hasan. The methodology to implement RHCOC algorithm done by Mohammad Nazmol Hasan and Md. Nurul Haque Mollah. The R package of the RHCOC algorithm developed by Md. Bahadur Badsha. The final validation of paper done by Mohammad Nazmol Hasan, Md. Bahadur Badsha and Md. Nurul Haque Mollah. The whole process was monitored by Md. Nurul Haque Mollah.

