## Supplementary material for "Robust Hierarchical Co-clustering to Explore Toxicogenomic Biomarkers and Their Regulatory Doses of Chemical Compounds": https://bsmrau.edu.bd/nazmol/other-activities/

**Contents**


**S1. Hierarchical Clustering Algorithm**

**S1.1.** Distance Measures

**S1.2.** Linkage or Clustering Methods

**S2. Data Contamination Models**

**S2.1.** Tukey-Huber Contamination Model (THCM)

**S2.2.** Independent Contamination Model (ICM)

**S3. Control Chart for Individual Measurement (CCIM)**


**Tables**


**Table S1:** Distance measures/methods considered for the RHCOC algorithm.

**Table S2.** The gene and DCCs cluster members of PPAR signaling pathway data retrieve by the RHCOC co-clustering algorithm.

**Table S3.** Functional annotation of KEGG pathway on the biomarker genes (genes in biomarker co-cluster 1) explored by the **RHCOC** algorithm for PPAR signaling pathway datasets.


**Figures**


**Figure S1.** Dendrogram of the row/gene and column/DCC cluster generated using the hierarchical clustering. In the figure **(A)** represents the row/gene clusters and **(B)** represents column/DCC clusters. There are four row/gene clusters and three column/DCC clusters are observed from the figure. These cluster numbers will be used as the input in the RHCOC algorithm for making co-clusters.

**Figure S2.** Hierarchical clustering using the distance and HC method combination manhattan: ward.D. **(A)** Gene clustering of GMP data. **(B)** Gene clustering of PPAR-SP data. **(C)** DCCs clustering of GMP data. **(D)** DCCs clustering of PPAR-SP data.

**Figure S3.** Average percentage of error rate is plotted against the combination of distance and HC methods when the dataset are simulated 100 times. **(A)** Average gene clustering ER in case of THCM data contamination method. **(B)** Average gene clustering ER in case of ICM data contamination method. **(C)** Average DCCs clustering ER in case of THCM data contamination method. **(D)** Average DCCs clustering ER in case of ICM data contamination method.

**Figure S4.** Co-cluster graph of the simulated dataset. In the graph row/gene clusters are plotted along the X-axis and the column/DCCs clusters are plotted along the Y-axis. Each row cluster in the X-axis make co-cluster with column/DCCs cluster in the Y-axis.

**Figure S5.** Control chart for individual measurement. The co-clusters mean are considered as the individual measurement. The co-clusters (a pair of row/gene and column/DCCs cluster makes a co-cluster) which mean plot beyond the UCL or LCL are considered as the biomarker co-clusters. In the figure **(A)** represents the simulated dataset, **(B)** represents glutathione metabolism pathway (GMP) dataset and **(C)** represents PPAR signaling pathway (PPAR-SP) dataset.

### S1. Hierarchical Clustering Algorithm

Hierarchical clustering (HC) algorithm clusters row or column entities of a data matrix based on different distance and linkage/clustering methods. The distance method determines how the distance between two observations is calculated. The linkage/clustering method is used to merge the row or column entities into clusters of data matrix based on the distance matrix. In the analysis of biological data the most commonly used clustering methods are of two types: Hierarchical and non-hierarchical (also known as partitioning). The hierarchical clustering approach builds clusters by repeatedly joining and merging the objects separated by the shortest distance. Following merging of the closest two points the distance matrix is updated and the process repeated until all objects are joined. The robust hierarchical co-clustering (RHCOC) algorithm five distance methods (euclidean, maximum, manhattan, canberra, minkowski) and seven HC clustering methods (single, complete, average, ward.D, ward.D2, mcquitty, median, and centroid) are considered. The distance and linkage or clustering methods are described below.

#### S1.1. Distance Measures

The distance measure quantifies the distance or dissimilarity among m-dimensional objects or items of a $n \times m$ data matrix. Actually this method determines how the distance between two observations is calculated. In this study we consider $n \times m$ gene-DCCs toxicogenomic data matrix consisting of G = $(G_1, G_2, \ldots, G_n)$ genes and C = $(C_1, C_2, \ldots, C_m)$ dose of chemical compounds (DCCs)/treatment combinations. As mentioned earlier we transform the gene-DCCs data matrix for robust results from RHCOC algorithm. Each of the input of this transformed data matrix is termed as $Tr(G_i, C_j)$ for the convenience of further use. This represents the transformed FCGE (average over animal sample) value for the $i^{th}$gene under the $j^{th}$treatment combination. In our RHCOC algorithm we consider the following distance methods (given in table). These distance methods (Table 1) measures distances between genes or row entities over the DCCs or column entities. Similarly, we can also calculate distances between DCCs or column entities.

**Table S1:** Distance measures/methods considered for the RHCOC algorithm.

| Distance Method | Mathematical Formulation |
|---|---|
| **Euclidean** | $d_{G_i,G_{i'}} = \left(\sum_{j=1}^{m}\left(Tr(G_i, C_j) - Tr(G_{i'}, C_j)\right)^2\right)^{1/2}$ |
| **Minkowski** | $d_{G_i,G_{i'}} = \left(\sum_{j=1}^{m}\left\|Tr(G_i, C_j) - Tr(G_{i'}, C_j)\right\|^v\right)^{1/v}$ |
| **Manhattan** | $d_{G_i,G_{i'}} = \sum_{j=1}^{m}\left\|Tr(G_i, C_j) - Tr(G_{i'}, C_j)\right\|$ |
| **Canbera** | $d_{G_i,G_{i'}} = \sum_{j=1}^{m}\frac{\left\|Tr(G_i, C_j) - Tr(G_{i'}, C_j)\right\|}{Tr(G_i, C_j) + Tr(G_{i'}, C_j)}$ |
| **Maximum** | $d_{G_i,G_{i'}} = max_j\left\|Tr(G_i, C_j) - Tr(G_{i'}, C_j)\right\|$ |

### S1.2. Linkage or Clustering Methods

We have described below the linkage or clustering methods that can be used in our RHCOC algorithm.

#### Single Llinkage

The single linkage clustering algorithm clusters objects (genes or DCCs) of toxicogenomic data based on the distance or similarity between two pairs of genes/DCCs. At the starting, the smallest distance $D = \{d_{G_i,G_{i'}}\}$ will be found and merge the corresponding genes and form a cluster $(G_i G_{i'})$. In the next step, the distance between the clusters $(G_i G_{i'})$ and $G_{i''}$ are computed a by

$$d_{(G_i,G_{i'})G_{i''}} = \min\{d_{G_i,G_{i''}}, d_{G_{i'},G_{i''}}\}$$

to form the cluster $(G_i, G_{i'} G_{i''})$. This process continues until all genes merge into a single cluster.

#### Complete Linkage

In complete linkage HC algorithm two objects form a cluster together, when their distance is the largest. The general agglomerative algorithm starts finding the minimum entry $D = \{d_{G_i,G_{i'}}\}$ and merge corresponding genes, such as $G_i$ and $G_{i'}$, to get cluster $(G_i, G_{i'})$. In the next step cluster clusters $(G_i G_{i'})$ and $G_{i''}$ will be merged into a cluster $(G_i, G_{i'} G_{i''})$ based on their maximum distance which is computed as

$$d_{(G_i,G_{i'})G_{i''}} = \max\{d_{G_i,G_{i''}}, d_{G_{i'},G_{i''}}\}$$

This process continues until all genes merge into a single cluster.

**Average Linkage**

Average linkage treats the distance between two clusters as the average between all pairs of items where one member of a pair belongs to each cluster. We begin searching the distance matrix $D = \{d_{G_i,G_{i'}}\}$ to find the nearest genes, for example, $G_i$ and $G_{i'}$ these objects are merged to get the cluster $(G_iG_{i'})$. In the subsequent step, the distance between $(G_iG_{i'})$ and cluster $G_{i''}$ is obtained by

$$d_{(G_iG_{i'})G_{i''}} = \frac{\sum_i \sum_{i''} d_{ii''}}{N_{G_iG_{i'}} N_{G_{i''}}}$$

Where $d_{ii''}$is the distance between gene $i$ in cluster $(G_iG_{i'})$ and gene $i''$ in cluster $G_{i''}$ and $N_{G_iG_{i'}}$ and $N_{G_{i''}}$ are the number of genes in clusters $(G_iG_{i'})$ and $G_{i''}$ respectively.

**Centroid**

The centroid method involves in finding out the mean vector for each of the clusters and talking distance between two centroids. Initially each of the genes in is cluster then the distance between clusters $G_i$ and $G_{i'}$ is

$$D = d_{\{\overline{Tr(G_i,C_.)},\ \overline{Tr(G_{i'},C_.\ )}\}}$$

**Median**

The median HC method seeks the median of each of the clusters and measure the distance between two median points. The distance between the median of two clusters $G_i$ and $G_{i'}$ is

$$D = d_{\{Tr(G_{i'},C_{Med}\ ),Tr(G_{i'},C_{Med}\ )\}}$$

**Ward's Algorithm**

Ward's HC algorithm clusters objects based on minimizing 'loss of information' from joining two groups. This algorithm used error sum of squares (ESS) to measure the loss of information. Firstly, for a given cluster r, let $ESS_r$ be the sum of squared deviations of every item in the cluster from the cluster mean (centroid). If there are r clusters, define ESS as $ESS = ESS_1 + ESS_2 + \cdots + ESS_r$. At each step in the analysis, the union of every possible pair of clusters is considered, and the two clusters whose combination results in the smallest increase in ESS (minimum loss of information) are joined. Initially, each cluster consists of a single item, and, if there are N items, $ESS_r = 0, r = 1, 2, \cdots N,$ so $ESS = 0$.

Clusters for the column entity or DCCs for this study can also be computed.

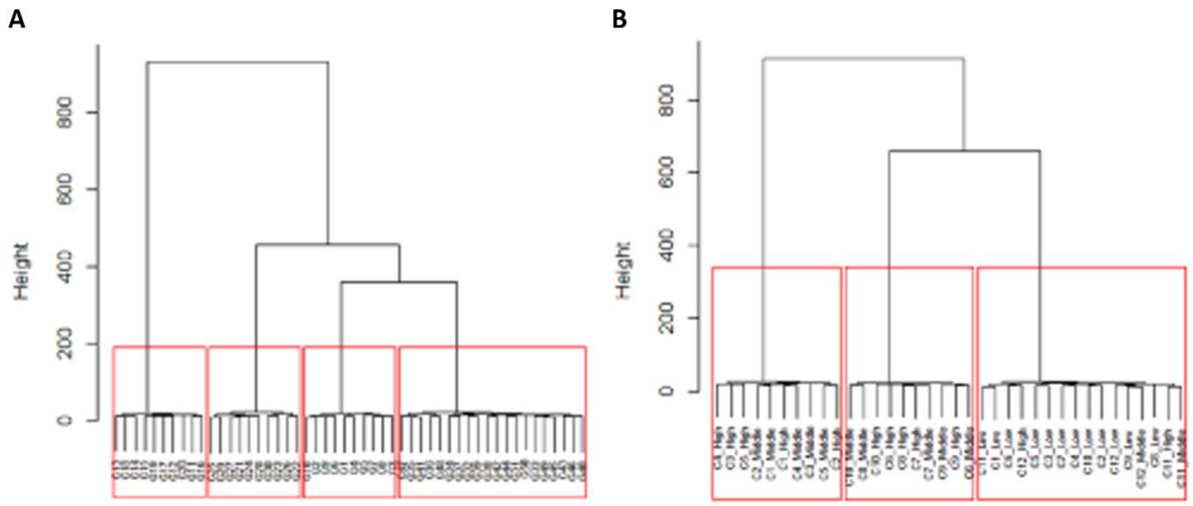


**Figure S1.** Dendrogram of the row/gene and column/DCC cluster generated using the hierarchical clustering. In the figure **(A)** represents the row/gene clusters and **(B)** represents column/DCC clusters. There are four row/gene clusters and three column/DCC clusters are observed from the figure. These cluster numbers will be used as the input in the RHCOC algorithm for making co-clusters.

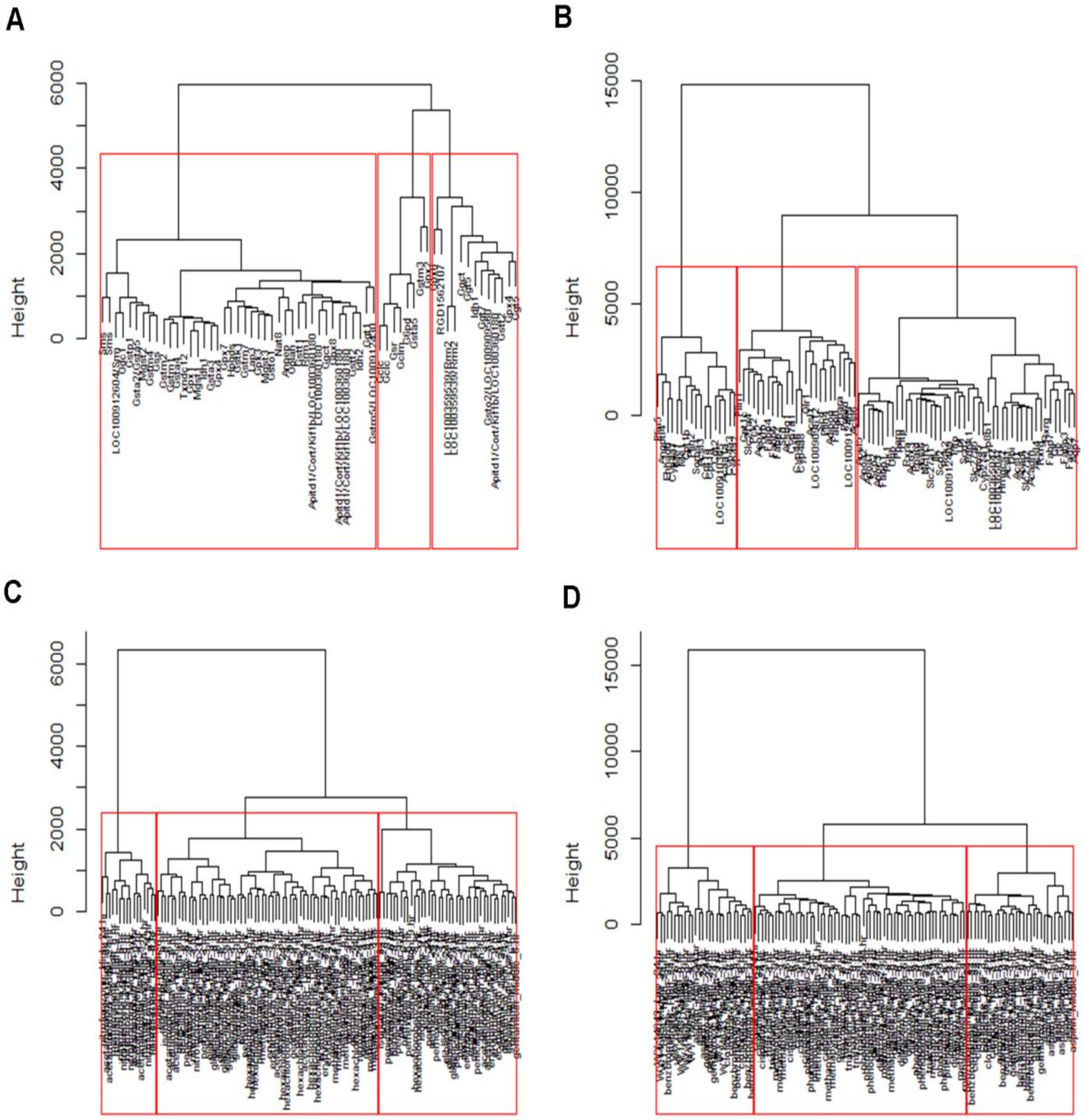


**Figure S2.** Hierarchical clustering using the distance and HC method combination manhattan: ward.D. **(A)** Gene clustering of GMP data. **(B)** Gene clustering of PPAR-SP data. **(C)** DCCs clustering of GMP data. **(D)** DCCs clustering of PPAR-SP data.

### S2. Data Contamination Models

The outlying observations in the dataset may arise casewise following the Tukey-Huber contamination model (THCM) (Agostinelli et al., 2015, Hasan et al., 2018) or independent cellwise following the independent contamination model (ICM) (Alqallaf et al., 2009, Hasan et al., 2018). The description of the THCM and ICM are given bellow.

#### S2.1. Tukey-Huber Contamination Model (THCM)

To examine the robustness of the proposed method we have contaminated the simulated gene-DCs FCGE dataset genewise/casewise using the following the THCM:

$$Z(D) = (1-\varepsilon)Z_0(D) + \varepsilon\tilde{Z}(D); \; 0 \le \varepsilon \le 0.5$$

Where, $\varepsilon$ is the small proportion of cases (genes) to be contaminated, $\tilde{Z}$ is the distribution of the FCGE values of the outlier contaminated genes, $Z_0$ is the nominal distribution by which we have generated the simulated data. In this study, to measure the robustness of the proposed algorithm we have considered wide range values of $\varepsilon$ (5%-40%).

#### S2.2. Independent Contamination Model (ICM)

We also have contaminated the simulated gene-DCs FCGE dataset following ICM:

$$D = (I - B_\varepsilon)D_{(0)} + B_\varepsilon\widetilde{D}$$

Where $D_{(0)} \sim Z_0$, $\widetilde{D} \sim \tilde{Z}$, $I$ is a $(m \times m)$ identity matrix, $B_\varepsilon = diag(B_1, B_2, \cdots, B_m)$and $B_j$ are independent $Birn(1, \varepsilon)$ it indicates that $\varepsilon$ is the probability of each component in $D$ to be contaminated. Additionally, the probability $\bar{\varepsilon}$ indicates that at least one component in $D$ to be contaminated. Where $\bar{\varepsilon} = 1 - (1 - \varepsilon)$. In this study we consider the value of $\bar{\varepsilon}$ are 0.000, 0.165, 0.304, 0.420, 0.517 and 0.598.

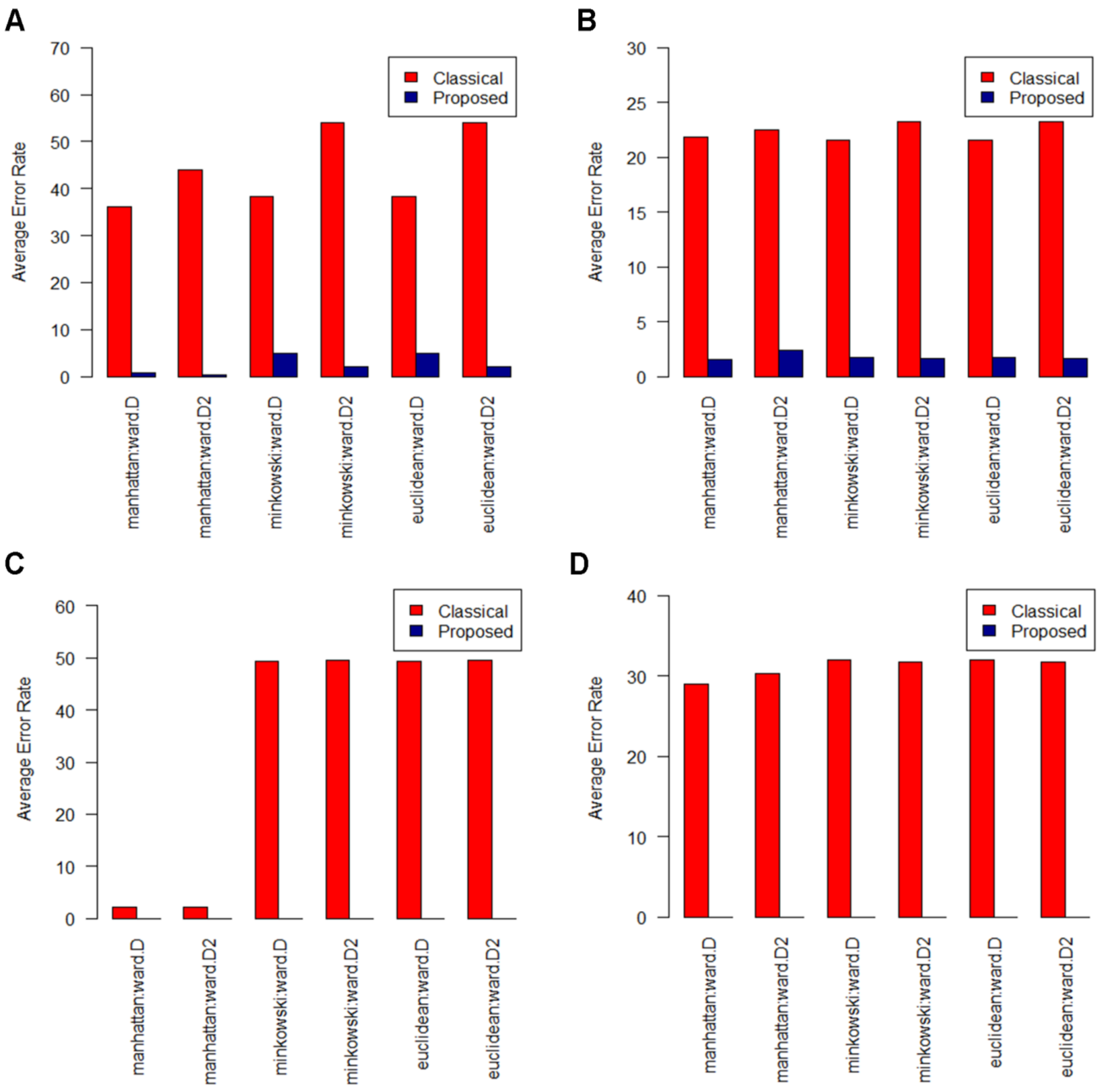


**Figure S3.** Average percentage of error rate is plotted against the combination of distance and HC methods when the dataset are simulated 100 times. **(A)** Average gene clustering ER in case of THCM data contamination method. **(B)** Average gene clustering ER in case of ICM data contamination method. **(C)** Average DCCs clustering ER in case of THCM data contamination method. **(D)** Average DCCs clustering ER in case of ICM data contamination method.

### S3. Control Chart for Individual Measurement (CCIM)

The control chart for individual measurement (CCIM) contains a central line (CL) which is the average value of the observations of quality characteristics, an upper control limit (UCL) and a lower control limit (LCL). CCIM or a control chart has a close connection with the hypothesis testing for drawing inferences. If an observation plots outside the control limits (UCL or LCL) it indicates that the process running with assignable causes of variation which is equivalent to rejecting the null hypothesis of statistical control and immediate action is necessary to eliminate that causes for quality output. Consequently, an observation plotting within the control limits is equivalent to accept the hypothesis of statistical control and no action is necessary for quality output (Montgomery, 2016). In fact, there is an analogy between the process monitoring consisting of individual measurement and assessing the DCCs/drugs toxicity based on toxicogenomic data (Hasan et al., 2019). We consider the each co-cluster mean of FCGE data as individual measurement to explore the biomarker co-cluster. Accordingly, any co-cluster mean falling outside the limit UCL or LCL we consider that co-cluster as the biomarker co-cluster and genes cluster and DCCs cluster that make this co-cluster are the clusters of toxicogenomic biomarkers and their regulatory DCCs. On the other side, the co-clusters which mean belong within the control limit (UCL and LCL) then the gene clusters and DCCs cluster which make these co-clusters are the clusters of non-biomarker genes and safe DCCs. In this regard we can calculate the CL, UCL and LCL in the following way.

Suppose there are $rk$ clusters in the gens or row entity and $ck$ clusters in the DCCs or column entity in the data matrix. Thus, according to robust hierarchical co-clustering (RHCOC) algorithm the gene (row) and DCCs (column) clusters together with make $rk \times ck = Z$ co-clusters. If we consider the $\bar{x}_z$ is the mean of the $z^{th}$ co-cluster then for our problem the observations/measurements are $\bar{x}_1, \bar{x}_2, \cdots, \bar{x}_z, \cdots, \bar{x}_Z$. Using these observations the CL, UCL and LCL can be calculated as:

$$MR_z = |\bar{x}_z - \bar{x}_{z-1}|$$

$$\overline{MR} = \sum_{2}^{Z} |\bar{x}_z - \bar{x}_{z-1}|$$

$$CL = \sum_{1}^{Z} \bar{x}_z = \bar{\bar{x}}$$

$$UCL = \bar{\bar{x}} + 3\frac{\overline{MR}}{d_2}$$

$$LCL = \bar{\bar{x}} - 3\frac{\overline{MR}}{d_2}$$

Where $MR_z$ is the moving range of two observations and if the moving range calculated based on two observations then $d_2 = 1.128$.

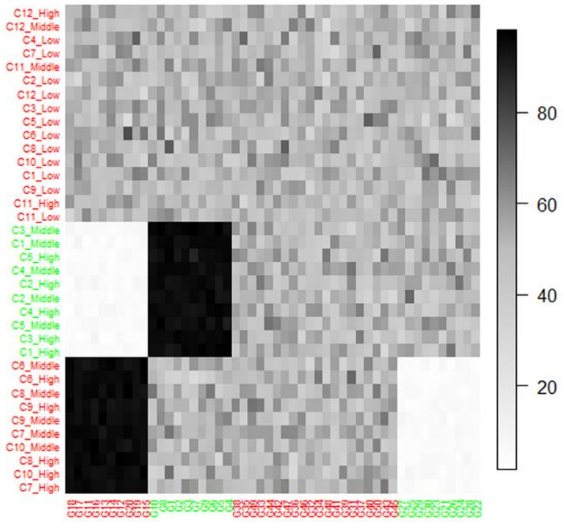


**Figure S4.** Co-cluster graph of the simulated dataset. In the graph row/gene clusters are plotted along the X-axis and the column/DCCs clusters are plotted along the Y-axis. Each row cluster in the X-axis make co-cluster with column/DCCs cluster in the Y-axis.

**Table S2.** The gene and DCCs cluster members of PPAR signaling pathway data retrieve by the RHCOC co-clustering algorithm.

| Cluster | Gene | DCC |
|---|---|---|
| **Cluster 1** | Cpt1a, Cyp4a3, Ehhadh, Plin5, LOC100910385, Cpt2, Acaa1a, Cyp4a1, Cpt1a, Cyp4a2, Angptl4, Cpt1b | aspirin_Low_24.hr, aspirin_High_24.hr, aspirin_Middle_24.hr, benzbromarone_Middle_6.hr, benzbromarone_High_9.hr, benzbromarone_Middle_9.hr, benzbromarone_High_24.hr, benzbromarone_High_6.hr, clofibrate_Middle_6.hr, clofibrate_High_24.hr, clofibrate_Middle_9.hr, clofibrate_High_6.hr, clofibrate_High_9.hr, gemfibrozil_High_24.hr, gemfibrozil_Middle_24.hr, gemfibrozil_Low_24.hr, WY.14643_High_6.hr, WY.14643_Middle_6.hr, WY.14643_Middle_24.hr, WY.14643_Low_3.hr, WY.14643_Low_24.hr, WY.14643_Middle_9.hr, WY.14643_Low_6.hr, WY.14643_High_9.hr, WY.14643_Middle_3.hr, WY.14643_High_3.hr, WY.14643_Low_9.hr, WY.14643_High_24.hr |
| **Cluster 2** | Rxra, Cyp8b1, Sorbs1, Plin2, Fabp2, Scp2, Pck2, Hmgcs2, Acsl4, LOC100912469, Ppard, Apoa5, Fabp3, Acox2, Me1, Acox3, Acsl5, Fads2, Slc27a5, Gk, Ilk, Slc27a2, Acsl5, Scp2, Rxrb, Pparg, Dbi, Lpl, Fabp7, | aspirin_High_3.hr, aspirin_Low_9.hr, aspirin_High_9.hr, aspirin_Low_3.hr, aspirin_Middle_9.hr, aspirin_Low_6.hr, aspirin_Middle_3.hr, aspirin_High_6.hr, aspirin_Middle_6.hr, benzbromarone_Low_3.hr, benzbromarone_High_3.hr, benzbromarone_Low_6.hr, benzbromarone_Middle_3.hr, benzbromarone_Low_9.hr, benzbromarone_Low_24.hr, benzbromarone_Middle_24.hr, clofibrate_Middle_3.hr, clofibrate_Middle_24.hr, clofibrate_High_3.hr, clofibrate_Low_6.hr, clofibrate_Low_3.hr, clofibrate_Low_9.hr, clofibrate_Low_24.hr, gemfibrozil_Low_9.hr, gemfibrozil_Middle_6.hr, gemfibrozil_High_9.hr, gemfibrozil_Low_6.hr, gemfibrozil_Middle_9.hr, gemfibrozil_High_3.hr, gemfibrozil_Middle_3.hr, gemfibrozil_High_6.hr |

| | | |
|---|---|---|
| | Acsl1, Pdpk1, LOC100365047, Apoc3, Apoa2, Gk, Pck1, Acsl1, Slc27a1, Scd2, Pltp, Acadl, Pltp, LOC100365047, Fabp1, Acsl3, Apoa1, Rxrg, Aqp7, Ubb, Rxrg, Acadm, Cyp27a1, Me1, Nr1h3 | |
| **Cluster 3** | Olr1, LOC100912469, Adipoq, Cyp4a8, Acsl6, Fabp5, Scd, Fabp6, Acsbg1, Ppard, Scd, Fabp4, Acsl1, Pck2, Ppara, Plin1, Plin4, Cpt1c, Cyp4a8, Cyp7a1, LOC100909612, Slc27a4 | cisplatin_Middle_6.hr, cisplatin_Low_24.hr, cisplatin_High_3.hr, cisplatin_Middle_24.hr, cisplatin_Middle_3.hr, cisplatin_High_6.hr, cisplatin_Low_6.hr, cisplatin_Low_3.hr, cisplatin_High_9.hr, cisplatin_Low_9.hr, cisplatin_High_24.hr, cisplatin_Middle_9.hr, diltiazem_High_6.hr, diltiazem_Middle_3.hr, diltiazem_High_24.hr, diltiazem_High_3.hr, diltiazem_High_9.hr, diltiazem_Middle_6.hr, diltiazem_Low_9.hr, diltiazem_Low_6.hr, diltiazem_Low_24.hr, diltiazem_Middle_24.hr, diltiazem_Low_3.hr, diltiazem_Middle_9.hr, gemfibrozil_Low_3.hr, methapyrilene_Middle_24.hr, methapyrilene_Low_9.hr, methapyrilene_High_6.hr, methapyrilene_Middle_3.hr, methapyrilene_High_24.hr, methapyrilene_High_3.hr, methapyrilene_Low_3.hr, methapyrilene_Low_6.hr, methapyrilene_High_9.hr, methapyrilene_Middle_9.hr, methapyrilene_Low_24.hr, methapyrilene_Middle_6.hr, phenobarbital_Middle_3.hr, phenobarbital_Low_24.hr, phenobarbital_High_9.hr, phenobarbital_High_24.hr, phenobarbital_Middle_24.hr, phenobarbital_Middle_9.hr, phenobarbital_Low_9.hr, phenobarbital_Middle_6.hr, phenobarbital_Low_3.hr, phenobarbital_Low_6.hr, phenobarbital_High_6.hr, phenobarbital_High_3.hr, triazolam_High_24.hr, triazolam_High_9.hr, triazolam_Low_6.hr, triazolam_Middle_6.hr, triazolam_Middle_9.hr, triazolam_Low_24.hr, triazolam_Low_3.hr, triazolam_High_3.hr, triazolam_Middle_24.hr, triazolam_High_6.hr, triazolam_Middle_3.hr, triazolam_Low_9.hr |

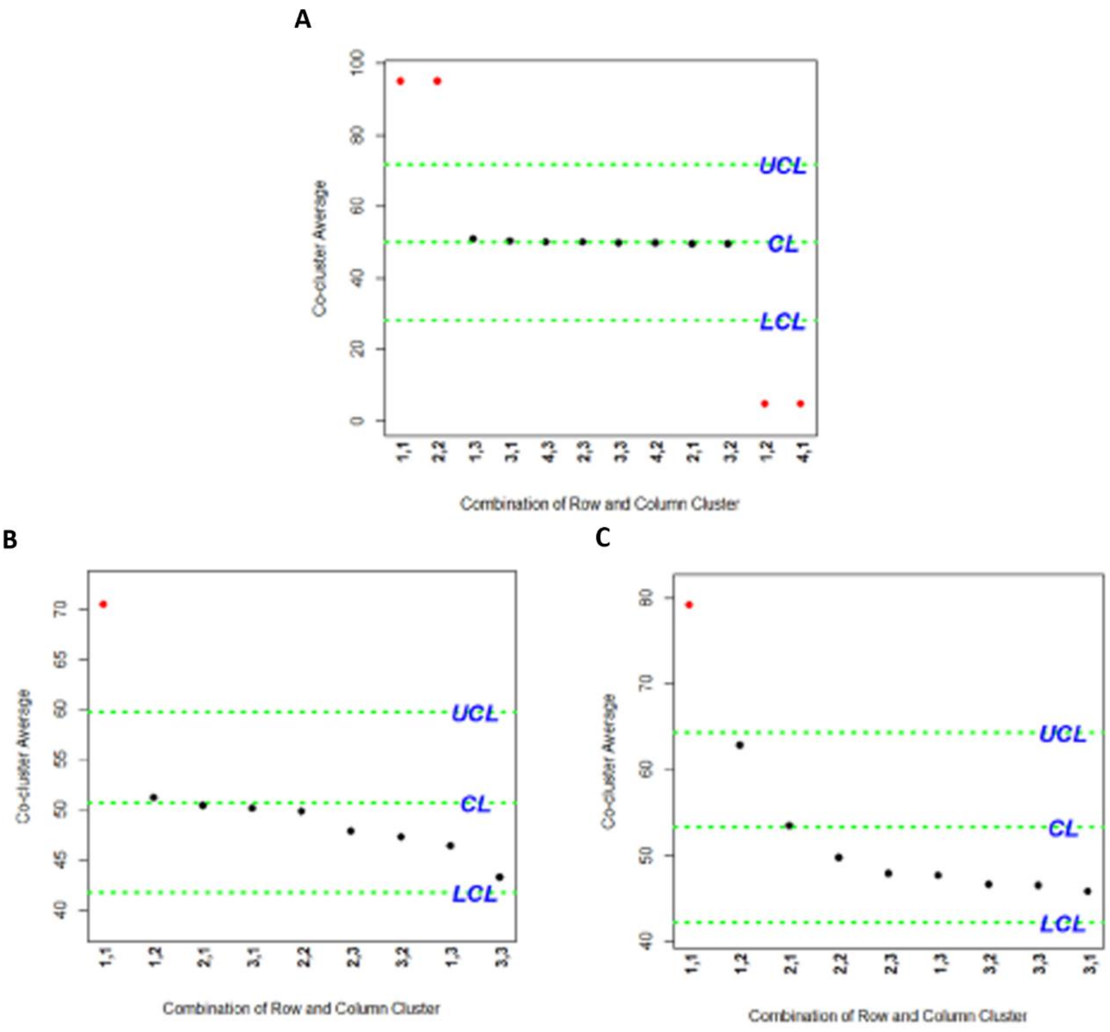


**Figure S5.** Control chart for individual measurement. The co-clusters mean are considered as the individual measurement. The co-clusters (a pair of row/gene and column/DCCs cluster makes a co-cluster) which mean plot beyond the UCL or LCL are considered as the biomarker co-clusters. In the figure **(A)** represents the simulated dataset, **(B)** represents glutathione metabolism pathway (GMP) dataset and **(C)** represents PPAR signaling pathway (PPAR-SP) dataset.

**Table S3.** Functional annotation of KEGG pathway on the biomarker genes (genes in biomarker co-cluster 1) explored by the **RHCOC** algorithm for PPAR signaling pathway datasets.

| | **Term** | **Count** | **%** | **p-value** | **Genes** |
|---|---|---|---|---|---|
| PPAR Signaling Pathway | rno03320:PPAR signaling pathway | 11 | 80.33 | 5.59E-20 | Sorbs1, Cpt1b, Cpt2, Ehhadh, Acsl3, Acaa1a, Cyp4a2, Cpt1a, Cyp4a3, Cpt1a, Angptl4, Cyp4a1 |
| | rno00071:Fatty acid degradation | 9 | 60.0 | 1.59E-16 | Cpt1b, Cpt2, Ehhadh, Acsl3, Acaa1a, Cyp4a2, Cpt1a, Cyp4a3, Cpt1a, Cyp4a1 |
| | rno01212:Fatty acid metabolism | 6 | 40.0 | 6.08E-9 | Cpt1b, Cpt2, Ehhadh, Acsl3, Acaa1a, Cpt1a, Cpt1a |
| | rno01100:Metabolic pathways | 7 | 46.66 | 0.004 | Me1, Me1, Ehhadh, Acsl3, Acaa1a, Cyp4a2, Cyp4a3, Cyp4a1 |
| | rno04920:Adipocytokine signaling pathway | 3 | 20.0 | 0.005 | Cpt1b, Acsl3, Cpt1a, Cpt1a |
| | rno00590:Arachidonic acid metabolism | 3 | 20.0 | 0.005 | Cyp4a2, Cyp4a3, Cyp4a1 |
| | rno00830:Retinol metabolism | 3 | 20.0 | 0.006 | Cyp4a2, Cyp4a3, Cyp4a1 |
| | rno04146:Peroxisome | 3 | 20.0 | 0.006 | Ehhadh, Acsl3, Acaa1a |
| | rno04750:Inflammatory mediator regulation of TRP channels | 3 | 20.0 | 0.011 | Cyp4a2, Cyp4a3, Cyp4a1 |
| | rno04270:Vascular smooth muscle contraction | 3 | 20.0 | 0.012 | Cyp4a2, Cyp4a3, Cyp4a1 |
| | rno00280:Valine, leucine and isoleucine degradation | 2 | 13.33 | 0.075 | Ehhadh, Acaa1a |
